# Towards a universal foundation model for biobank-scale human genome variation

**DOI:** 10.1101/2025.01.29.635579

**Authors:** Augix Guohua Xu, Yu Xu, Yiming Xing, Pengchao Luo, Jianbo Yang, Yinqi Bai, Kun Tang

**Author notes:** Corresponding authors: Kun Tang, Yinqi Bai, Jianbo Yang.

## Abstract

Millions of human genomes have been genotyped by national biobanks worldwide. Training large language models (LLM) with this data may lead to a universal model of human genome with tremendous potential. Yet the quadrillions (10^15^) of nucleotides— resulting from genome length multiplied by population size—pose formidable challenges for modeling. In this study, we propose a novel AI framework designed to scale with this data and support diverse analytical tasks. To demonstrate this scheme, we developed SNPBag—a foundation model focusing on single nucleotide polymorphism (SNP). With 0.8 billion parameters, it is trained on one million synthesized human genomes, corresponding to a total of 6 trillion SNP tokens. SNPBag showed superior performance in benchmarking of multiple tasks. In genotype imputation, it achieves state-of-the-art (SOTA) accuracy. In haplotype phasing, it rivals the best method with a 72-fold speedup. By encoding 6 million SNPs per genome into a 0.75 MB embedding, SNPBag enables efficient storage, transfer and downstream applications. In particular, the genome embeddings facilitate rapid ancestry inference across global populations and detection of genetic relationships up to 12th-degree relatives. Collectively, SNPBag introduces a new paradigm for scalable, unified and multitask analysis of the ever-growing human variation data.

## Introduction

Artificial intelligence, especially foundation models based on transformers, has revolutionized biomedical research, enabled comprehensive omics-scale analyses and reshaped our understanding of complex biological systems. In proteomics, the AlphaFold series accurately predicts protein structures and accelerates drug discovery (*1*). In transcriptomics, particularly single-cell transcriptomics, models including Geneformer, scGPT, and scBERT support cell type annotation, gene regulatory network inference, and perturbation response prediction (*2*–*4*). In genomics, models including Nucleotide Transformer (*5*) and EVO (*6*) process full genome sequences across many species to predict gene essentiality and annotate genomic elements.

In contrast, for single-species genome data—such as human genomes—there remains a lack of AI foundation models capable of handling the vast genomic variations among individuals. This gap is striking, given both the sheer volume and critical significance of human genomic data. Over the past two decades, human genome data has expanded exponentially. Landmark initiatives such as the International HapMap Project (*7*) and the 1000 Genomes Project (*8*), systematically examined thousands of human genomes and catalogued global genetic variation. Large-scale biobanks of “hundreds of thousands to millions of individuals”—including the UK Biobank (*9*), the Million Veteran Program and the All of Us Research Program (*10*), the China Kadoorie Biobank (*11*), Biobank Japan (*12*), deCODE Genetics (*13*), the Estonian Biobank (*14*), FinnGen (*15*), and Lifelines (*16*)—provide extensive population-scale genomic datasets. Most recently, The Human Genome Project II initiative proposes sequencing up to 1% of the global population, or 80 million individuals (*17*).

Human genome data presents unique challenges for AI model design. The human genome is extremely large (~3 billion nucleotides), and biobanks now contain millions of individual genomes. Designing models to process full genome sequences— following frameworks like current DNA language models such as EVO-2—would require learning quadrillions of tokens (10^15^ ~ 10^16^), far exceeding even the 10^12^ ~ 10^13^ token used to train the most advanced large language models (*18, 19*), making this approach infeasible in the coming years. Furthermore, human genomes are nearly identical, differing by only 0.1~0.6% between individuals (*20*), rendering full-sequence modeling unnecessarily redundant.

In this study, we propose a framework with a large language foundation model designed for the universal processing of biobank-scale human genome data, defined by four key features:

1. Variant-centered tokenization that focuses on polymorphic sites rather than every nucleotide, dramatically reducing computation.
2. Whole-genome modeling that analyzes the entire genome rather than partial genomic windows, capturing long-range linkage disequilibrium, extended haplotypes, and—most importantly—genome-wide epistatic interactions underlying human phenotypes.
3. Compact genome embeddings that provide low-dimensional representations, enabling the development of downstream applications even with limited computational resources and modest sample sizes.
4. Multitask training that unifies a broad spectrum of human genomic variation analyses within a single framework.

To explore the feasibility of this framework, we developed SNPBag—a novel foundation model specialized in processing whole-genome single nucleotide polymorphisms (SNPs) data. SNPs are the most prevalent form of genetic variation, making them ideal for large-scale modeling (*7*). Furthermore, SNPs also serve as the primary substrate for genome-wide association studies (GWAS)—a core biobank application linking genotypes to phenotypes. To date, over 110,000 GWAS studies have been conducted, covering approximately 15,000 traits and millions of genomes (*21*). Beyond GWAS, common SNP analyses include genotype imputation, haplotype phasing, population stratification, and relatedness inference. Leading bioinformatic tools include IMPUTE, BEAGLE, MINIMAC, and EAGLE for imputation (*22*–*25*); BEAGLE, SHAPEIT, and EAGLE for haplotype phasing (*22, 24*–*26*); PLINK PCA and nonlinear methods (t-SNE, UMAP) for population stratification (*27*–*29*); and KING, ERSA, PRIMUS, and Bonsai for relatedness inference (*30*–*32*). While effective, these tools face limitations in scalability, computational efficiency, reference panel dependence, and task-specific fragmentation. Recently, neural network approaches have also emerged (*33*–*39*). However, these methods primarily provide end-to-end solutions for specific tasks and share similar limitations with conventional tools, complicating the selection of optimal methods.

Here, we present SNPBag, the first foundation model designed for large-scale human SNP data. Training on 1 million genomes, 6 million SNP sites and 6 trillion tokens, SNPBag internalized complex SNP structures, enabling broad-range tasks, benchmarked in genotype imputation, haplotype phasing, genome embedding, population classification, and relatedness inference. When compared against current state-of-the-art methods, SNPBag demonstrated exceptional performance across these tasks. we have made the model inference publicly available on GitHub (https://github.com/augix/SNPBag) and invite the research community to explore additional application scenarios.

## Results

### Base model pre-training

SNPBag, a model built on Bidirectional Encoder Representations from Transformers (BERT), leverages an attention mechanism to capture complex dependencies among SNP sites within long genomic distances, thereby generating contextualized genotype representations. These representations support the decoding of diverse features for individual genomes, as outlined in the SNPBag workflow (Fig. 1a). To address privacy issue and increase sample size, we simulated 1 million phased genomes using publicly available data (Fig. 1a, top). These synthetic genomes were used to pre-train a base model and subsequently fine-tune task-specific models (vertical arrows, Fig. 1a). During inference (horizontal arrows, Fig. 1a), input data in various formats were processed as sequences by trained models to perform tasks such as genotype imputation, haplotype phasing, and genome embedding generation. The resulting compressed genome embeddings enabled downstream applications, including population classification, relatedness inference, and potentially phenotype prediction.

**Fig. 1:**
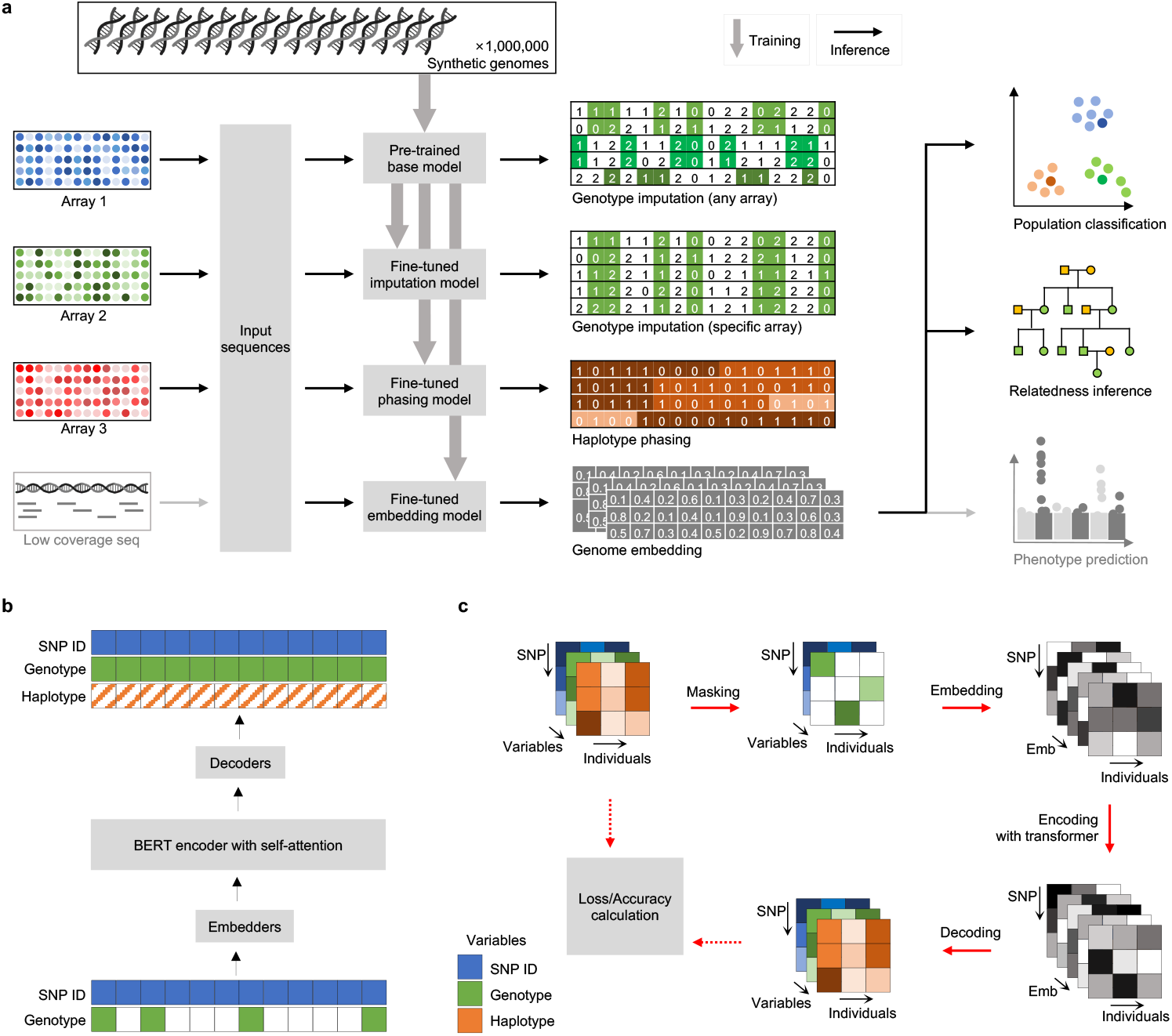
Model architecture. **a**, Overview of SNPBag. One million synthetic genomes (top) were used for pre-training and fine-tuning across tasks (vertical arrows). The base model (also in panel b), pre-trained via masked genotype prediction, functions directly as a genotype imputer for any SNP arrays, and potentially for low-coverage sequencing data. Fine-tuning the base model enhances imputation accuracy for specific SNP arrays. Haplotype phasing is achieved by fine-tuning the base model with a new decoding head to predict haplotype switch states. Additionally, training an embedding head enables SNPBag to compress genome-wide SNPs into low-dimensional genome embeddings, supporting downstream tasks such as population classification and relatedness inference. **b**, The base model adopts a BERT architecture. During pre-training, portions of genotype sequence values (green squares) are masked (white squares), combined with their corresponding SNP IDs, and encoded by the model. Genotype values were decoded in the output. This structure is also adapted for fine-tuning of haplotype phasing (brown) by training a phase decoder. **c**, Detailed model architecture. In a single batch of input data with three dimensions, the SNP IDs and masked genotypes are combined into the variables dimension, and converted to an “Emb” dimension to match the attention-based encoder. During decoding, either masked genotypes (green) or haplotype phases (brown) are recovered from model output and compared to original values to compute loss and accuracy.

For pre-training (Fig. 1b, c), we masked 85–99% of genotypes randomly and paired them with corresponding SNP identifiers to form input sequences. These sequences were projected into a 512-dimensional space using linear embedding layers, feeding into a BERT encoder with 16 transformer layers, each with 16 attention heads. The encoder output was processed by a multilayer perceptron (MLP) to predict genotype values at masked positions, with model weights optimized using cross-entropy loss between predicted and actual genotypes. By swapping decoders, the base model was fine-tuned for tasks such as haplotype prediction (brown squares, Fig. 1b, c). Using this architecture, we trained both pilot and full models on distinct reference datasets. A chromosome 22 (chr22) model with 38 million parameters was developed for benchmarking imputation and phasing, while a whole-genome model with 836 million parameters was trained for demonstrating more applications. To showcase transfer learning, the chr22 model was pre-trained on 1 million individuals synthesized from the 1000 Genomes Project (1kG) samples, and evaluated directly on Human Genome Diversity Project (HGDP) samples. The synthetic data, generated by software RESHAPE (*40*), is referred to as the RESHAPE data henceforth. The whole-genome model, designed for public use, was pre-trained on 1 million simulated genomes of the HAPNEST project (*41*), which synthesized from both 1kG and HGDP samples to maximize genetic diversity.

SNP sites were restricted to single-nucleotide polymorphisms shared between 1kG and HGDP datasets with a minor allele frequency (MAF) >1% (as determined from 1kG data). The whole-genome model was trained on 6,030,514 SNPs, organized into consecutive sequences of 81,920 sites each (analogous to genomic contigs and hereafter referred to as “contigs”), whereas the chr22 model used a single 81,920-site contig, covering ~92% of chromosome 22.

### Genotype imputation

Genotype imputation infers untyped variants, enhancing data completeness for GWAS, reducing sequencing depth requirements, and harmonizing diverse genotyping platforms for broader analyses. The imputation relies on the linkage disequilibrium (LD) structure, where adjacent SNPs tend to be inherited together. To benchmark SNPBag, we evaluated imputation performance on 216 HGDP individuals genotyped with the Illumina Omni2.5 array on chromosome 22, masking non-array SNPs to impute the full SNP set. Traditional pipelines combining phasing software (SHAPEIT4, BEAGLE5.2, EAGLE2) and imputation software (IMPUTE5, MINIMAC4, BEAGLE5.2), with or without the 1KG reference panel, were used for comparison (*22*–*25, 42*).

The SNPBag base model achieved 96.78% imputation accuracy (concordance rate), ranking 11th among all the 20 tests in comparison (Fig. 2a). To optimize imputation performance, we fine-tuned the base model with the Omni2.5-specific masking pattern rather than random masking, generating the SNPBag omni model. Scaling-law experiments across training set sizes of 1 million, 100 thousand, 10 thousand, and 2 thousand individuals revealed diminishing accuracy gains with smaller datasets, with over-fitting evident below 100 thousand samples (Supplementary Fig. 1a, b). Fine-tuning on 1 million synthetic individuals (the same RESHAPE data used in pre-training) yielded state-of-the-art (SOTA) accuracy of 97.16% after approximately one epoch, surpassing the best traditional pipeline (BEAGLE5.2+IMPUTE5+reference panel, 97.10%) with statistical significance (paired t-test, *p =* 2.7×10^−4^, Fig. 2a). Across minor allele frequency (MAF) intervals with comparable SNP counts, the fine-tuned SNPBag omni model consistently outperformed other methods, exceeded even the average accuracy of reference-based methods (REF) by 0.1–0.7% across intervals (Fig. 2b). It also achieved the highest accuracy across all continental groups (red boxes in Fig. 2c). The population-specific accuracy was highest in Americans (AMR, 98.3%), remained relatively high among all the out-of-African (OOA) groups (96.2 - 97.8%), but was lower in Africans (AFR, 94.2%) with greater inter-individual variance (Fig. 2c, Supplementary Fig. 2). The reduction in AFR accuracy reflects the higher genetic diversity and weaker inter-site LD, both of which increase imputation uncertainty in prediction (*20*).

**Fig. 2:**
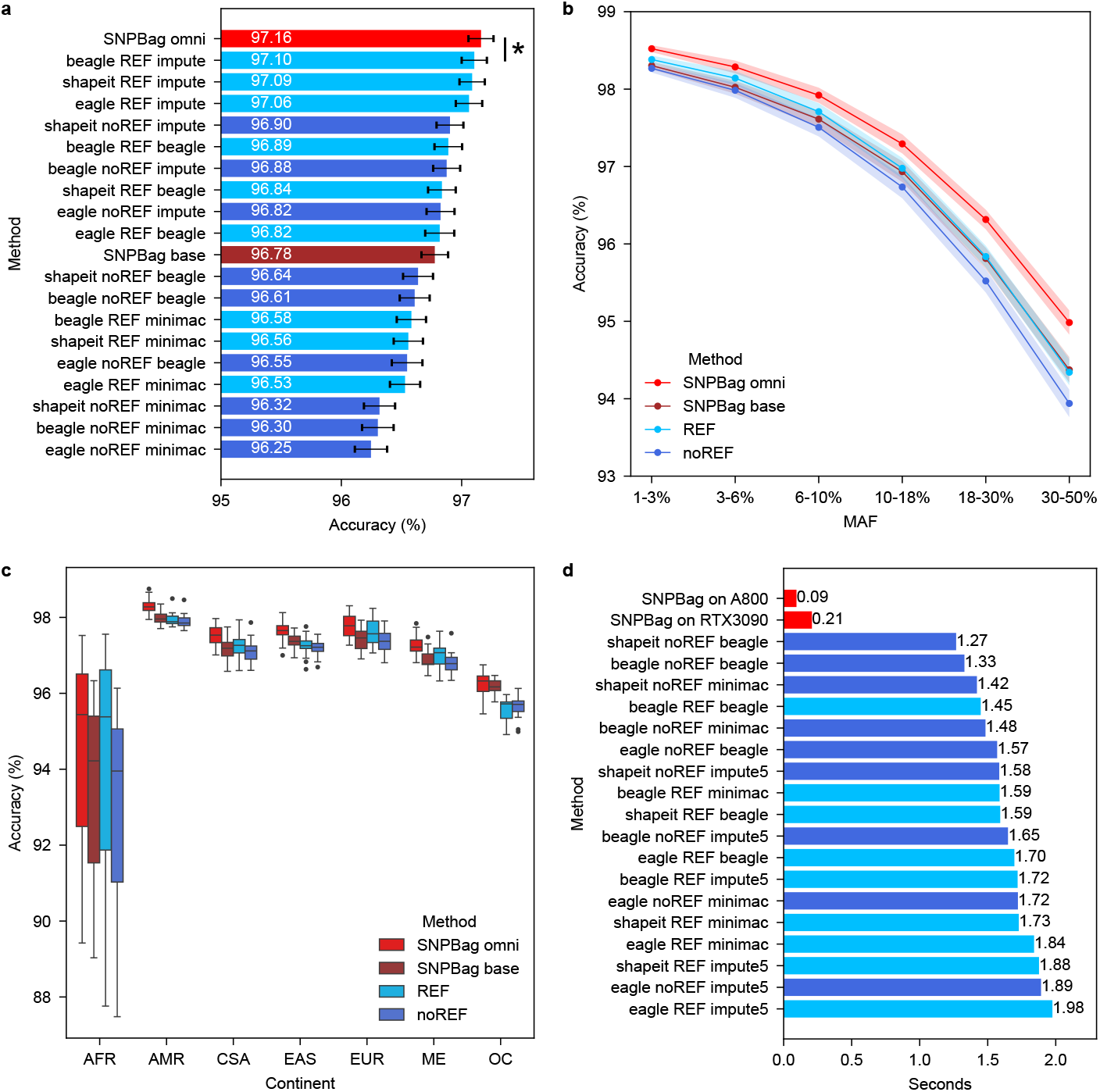
Genotype imputation benchmarking. **a**,**d**, Performance comparisons of SNPBag fine-tuned model (red) and base model (brown) versus traditional methods with reference panel (light blue), or without reference panel (dark blue) for imputation accuracy (a) and inference speed (d). The star mark (*) indicates statistical significance *p-value* < 0.05. A800 and RTX3090 are Nvidia chips. **b**, Imputation accuracy of different methods across bins of minor allele frequency, with roughly same numbers of SNPs within each bin. REF and noREF represent the average of traditional methods with and without the reference panel. **c**, Boxplot of imputation accuracy by methods and continental groups. Continental groups included Africans (AFR), Americans (AMR), Central and South Asians (CSA), East Asians (EAS), Europeans (EUR), Middle East populations (ME) and Oceanians (OC). Standard errors of mean estimations were plotted as error bars in panel a and as shadows in panel b.

Notably, SNPBag achieved an average inference time of ~0.09 seconds on chromosome 22 using one NVIDIA A800 GPU, representing a 13 to 20-fold speedup compared with traditional methods (on a single Intel® Xeon® Platinum 8458P CPU, Fig. 2d). Overall, SNPBag’s reference-free imputation and flexibility across arbitrary genotyping arrays highlight its potential as an efficient and scalable tool for genomic analysis.

### Haplotype phasing

Haplotype phasing separates alleles into maternal and paternal chromosomes, generating distinct haplotypes that improve disease association detection and clarify inheritance patterns. These haplotype blocks are fundamental to the diploid human genome. SNPBag’s base model, pre-trained to capture haplotype distributions, was fine-tuned for phasing across the full SNP set using a new MLP head to predict switch states between consecutive heterozygous sites. We trained the model on 1 million phased genomes (the same RESHAPE data used in pre-training and imputation fine-tuning), and tested on 26 physically phased HGDP individuals on chromosome 22. Scaling-law experiments revealed that smaller training sets (2,000 and 10,000 compared to 1 million) reduced accuracy and led to earlier over-fitting (Supplementary Fig. 1c, d).

With 1 million individuals, SNPBag achieved a 3.07% switch error rate, outperforming traditional reference-free methods (6.27-7.04%) but falling slightly short of the methods using the reference panel (2.54-2.70%, Fig. 3a). This performance gap likely reflects the limitation of training on “true phases” derived from traditional phasing outputs (*20*), which constrain the attainable accuracy. Across continental groups, SNPBag’s performance closely matched the average of reference-panel-based methods, with highest errors in Africans, followed by Oceanians, and generally low errors in other groups, respectively (Fig. 3b). Elevated African error rates (5.0% in SNPBag and 3.9% in reference-panel-based methods, respectively) likely arise from higher haplotype diversity, whereas Oceanian phasing errors may reflect underrepresentation of their haplotypes in both training data and reference panels.

**Fig. 3:**
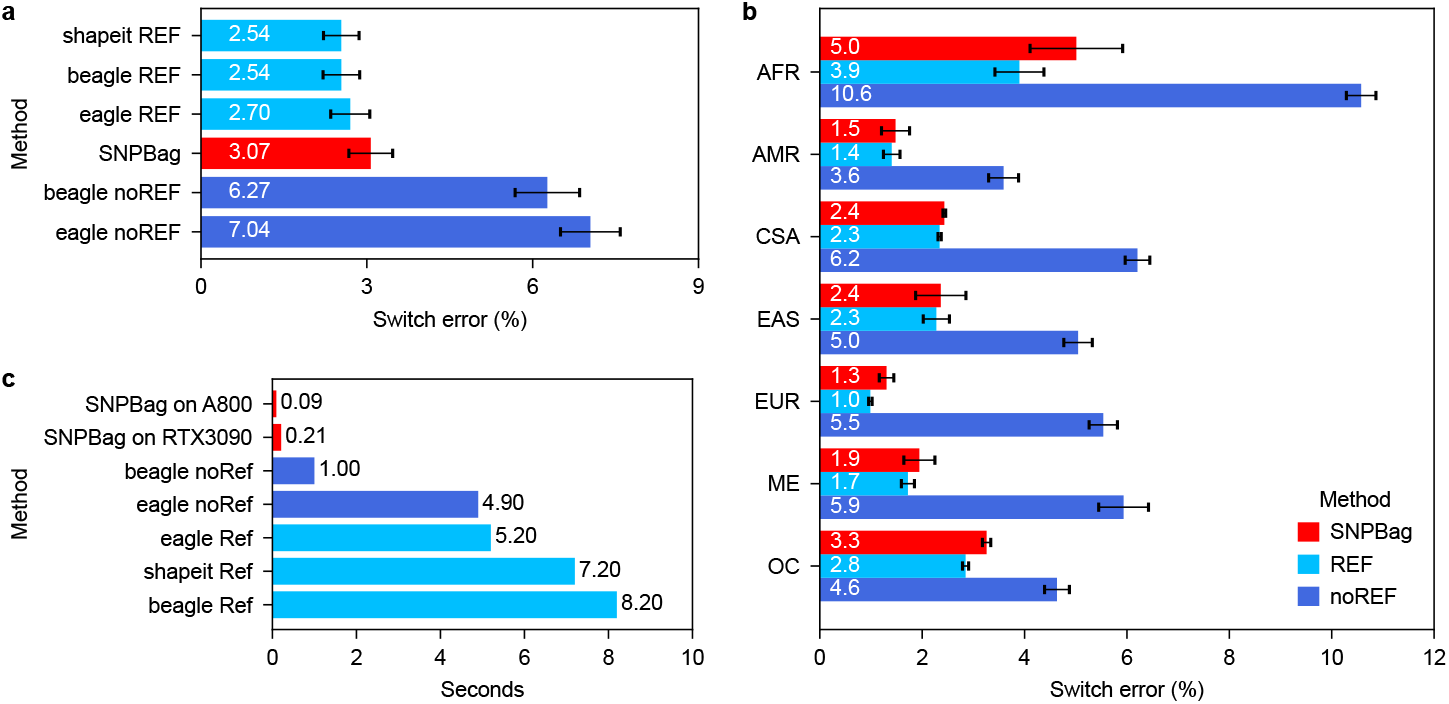
Haplotype phasing benchmarking. **a**, Comparisons of switch error rates between SNPBag (red) and traditional methods with reference panel (light blue) or without reference panel (dark blue). **b**, Phasing performance of different methods across 7 continental groups. **c**, Comparisons of inference speed between SNPBag (red) and traditional methods with reference panel (light blue) or without reference panel (dark blue). A800 and RTX3090 are Nvidia chips. Standard errors of mean estimations were plotted as error bars in panel a and b.

Notably, SNPBag’s reference-free phasing achieved ~0.09 seconds per sample on chromosome 22 using a single NVIDIA A800 GPU, delivering an 11 to 91-fold speedup over traditional methods (1.0–8.2 seconds on an Intel® Xeon® Platinum 8458P CPU, Fig. 3c). These results highlight SNPBag’s efficiency and flexibility, with future accuracy gains expected once training on experimentally determined haplotypes becomes feasible (*43*).

### Genome embedding

Whole-genome analyses, such as GWAS and population structure inference, pose computational challenges due to the vast number of loci and large cohort size. Representing genomes in low-dimensional spaces reduces complexity and enables efficient downstream applications. We first pre-trained SNPBag on genome-wide SNPs using 1 million HAPNEST-synthesized genomes derived from 1KG and HGDP panels (*41*). Pre-training was performed in parallel across all 22 autosomal chromosomes, completing 0.2 epoch (about 200,000 individuals), and yielded a base model that achieved 97.7% genotype imputation accuracy on the Omni2.5 array (Supplementary Fig. 3a).

For genome embedding, we divided the genome into 2,934 consecutive contigs, each containing 2,048 SNPs. Two sequential MLPs compressed each contig’s output into a 128-length embedding, with a third MLP decoder recovering the original contig sequence at 95% accuracy (Supplementary Fig. 3b), which can be improved with more computing resources. Notably, the resulting whole-genome embedding occupies only ~0.75 megabytes (751,104 bytes, calculated as 2,934 × 128 dimensions ×2 byte per dimension), enabling efficient storage, transfer, encryption, and large-scale analysis (Supplementary Fig. 3c). This compact representation enhances the scalability and practicality of genomic analyses, supporting diverse applications without compromising accuracy and privacy. Here, we demonstrate its application in population classification and relatedness inference.

### Population classification

Genome embeddings, which capture high-resolution SNP distributions, enable population structure analysis that is critical for GWAS correction and forensic ethnicity inference (*28, 44*). While conventional PCA-based methods have limited resolution at sub-continental groups, machine learning approaches are better suited to resolving fine-scale structures (*28, 29*). We generated embeddings for all 3,202 1KG samples and visualized them with UMAP, revealing distinct sub-continental clusters (Fig. 4a). In details, Japanese are separated from Chinese Han (CHB, CHS) and Dai-Kinh (CDX, KHV) clusters; Europeans split into Finnish, North-Western (CEU, GBR), and Southern (TSI, IBS) clusters; American groups (PUR, CLM, MXL, PEL) are clearly distinguishable; South Asian populations (PJL, STU, BEB) cluster closely but apart from Gujarati (GIH); and African populations (GWD, LWK, MSL) form distinct clusters, with some overlap between Yorubans (YRI) and Esans (ESN), and between African Caribbean (ACB) and African Americans (ASW) (Fig. 4a).

**Fig. 4:**
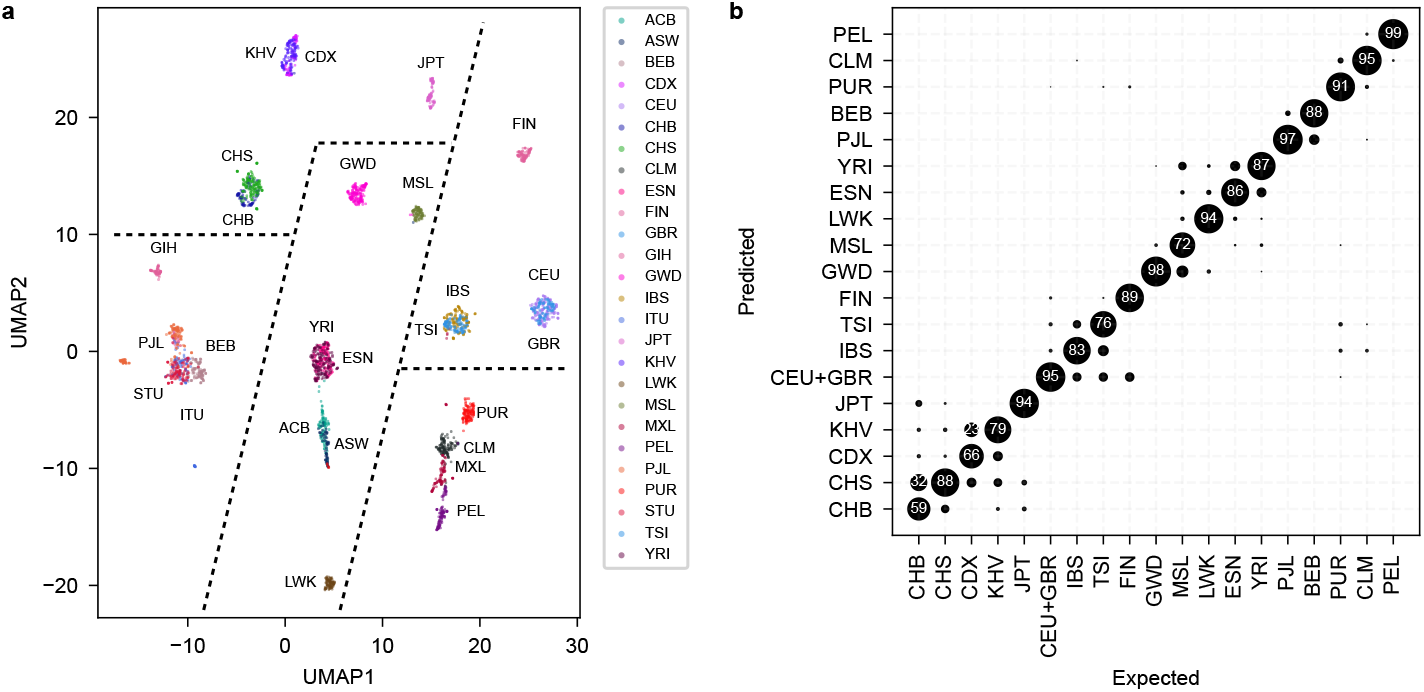
Population classification across 1000 Genomes Project (1KG) populations. **a**, UMAP visualization showing clustering of genome embeddings among 26 global populations, including 7 African (AFR) groups: Yoruba in Ibadan, Nigeria (YRI); Luhya in Webuye, Kenya (LWK); Mende in Sierra Leone (MSL); Esan in Nigeria (ESN); African Ancestry in Southwest USA (ASW); African Caribbean in Barbados (ACB); Gambian in The Gambia (GWD); 5 East Asian (EAS) groups: Han Chinese in Beijing (CHB); Japanese in Tokyo (JPT); Southern Han Chinese (CHS); Chinese Dai in Xishuangbanna (CDX); Kinh in Ho Chi Minh City, Vietnam (KHV); 5 European (EUR) groups: British in England and Scotland (GBR); Utah Residents with Northern and Western European Ancestry (CEU); Toscani in Italy (TSI); Finnish in Finland (FIN); Iberians in Spain (IBS); 5 South Asian (SAS) groups: Gujarati Indian in Houston (GIH); Punjabi from Lahore, Pakistan (PJL); Bengali in Bangladesh (BEB); Indian Telugu in the UK (ITU); Sri Lankan Tamil in the UK (STU); and 4 American (AMR) groups: Colombian in Medellín (CLM); Mexican Ancestry in Los Angeles (MXL); Puerto Rican in Puerto Rico (PUR); Peruvian in Lima (PEL). **b**, Confusion matrix of classification results for 19 populations, areas of circles represent percentages of individuals in expected classes. Big percentages (>20%) were shown inside circles.

By training a lightweight neural net with genome embeddings of all 1KG samples, we achieved an F1 score of 0.998 in classifying five continental super-populations (EUR, EAS, SAS, AMR, AFR) (Supplementary Fig. 4a). For the finer classification of 19 sub-populations, our model attained an average F1 score of 0.861, surpassing previously benchmarked SVM and GTM models (F1 = 0.8) (*29*), excelling in sub-continental discrimination (Fig. 4b). Results across all 26 populations were consistent with UMAP distributions (Supplementary Fig. 4b).

### Relatedness inference

Inferring relatedness is critical for case-control studies, where cryptic relatedness can introduce false associations (*31*), and for improving rare disease variant detection through shared ancestry (*45*). It also supports forensic and direct-to-consumer kinship analysis (*32*). Distant relatedness relies on detecting long identical-by-descent (IBD) haplotype segments (*46*).

Because SNPBag’s genome embeddings preserve extensive haplotype information, they should support robust inference of distant relatedness. To this end, we developed an MLP-based relatedness estimator using whole-genome embeddings. Trained on simulated deep pedigrees (up to 9th cousins) derived from 1KG founders, the estimator recalled relationships up to the 12th degree (Fig. 5a). At a detection power of 0.98-1.00, recall rates for exact matched degrees (D_0_) ranged from 0.96-1.00 for close relatives (degrees 1–2), 0.34–0.82 for intermediates (degrees 3–6), and 0.24-0.27 for distant relatives (degrees 7–9) (Fig. 5b). Exact match accuracy was low (0–0.09) beyond degree 10; however, allowing errors of 1–3 degrees (defined as D_1_, D_2_ and D_3_ respectively) yielded high accuracy across degrees 1 – 12, with the accuracy remaining above 0.75 up to the 8th, 10th and 11th degree relationships for D_1_, D_2_ and D_3_ respectively (Fig. 5b). Overall accuracy is comparable to established pairwise relatedness inference methods (*31*).

**Fig. 5:**
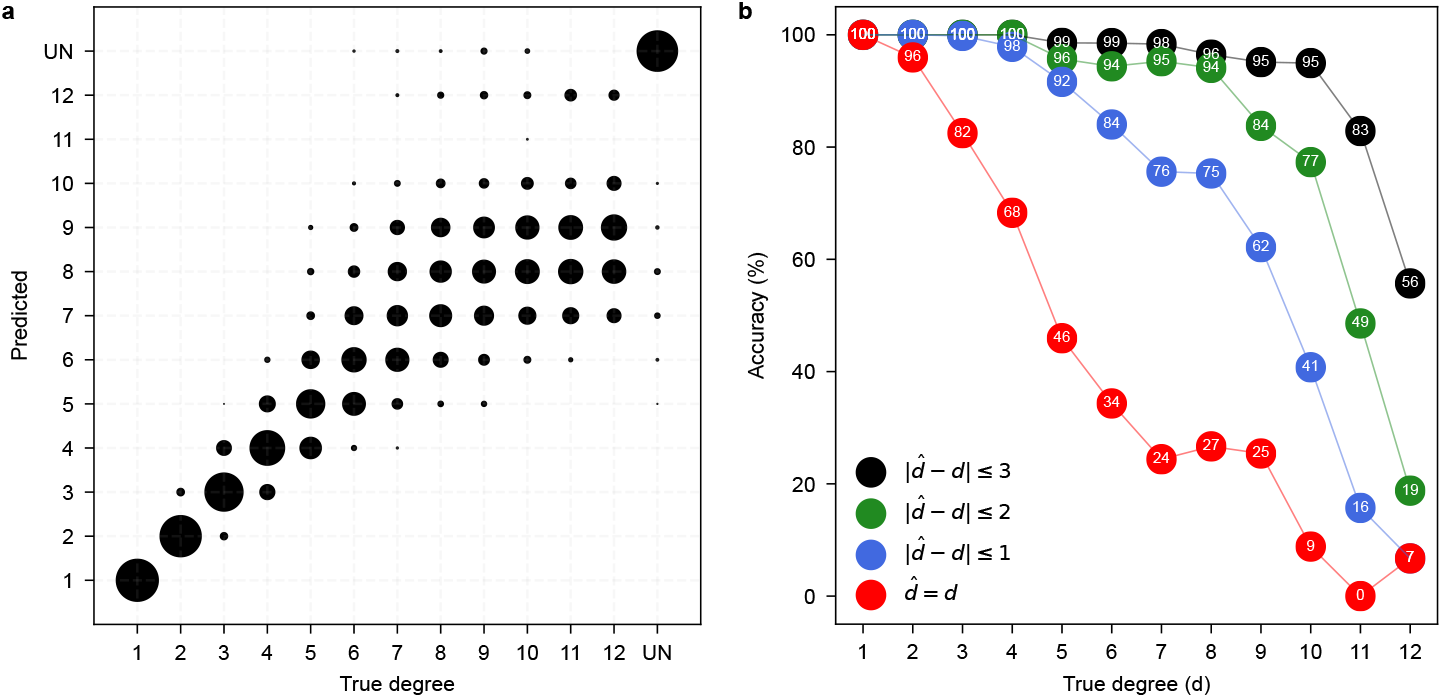
Performance of relatedness estimation. **a**, the confusion matrix of the estimated degree of relatedness against the true relationship. Area of the circles represent percentages of pairs in each degree of relatedness. UN: unrelated. **b**, the accuracy of inference was evaluated at four error range allowance: D_0_ in red: exact match between the estimation and truth; D_1_ in blue: 1-degree mismatch allowed; D_2_ in green: up to 2 degrees of mismatch allowed; and D_3_ in black: up to 3 degrees of mismatch allowed. The numbers in the curves indicate accuracy in percentage.

When benchmarked against the industry gold standard 23andMe’s Bonsai package (*32*), SNPBag’s accuracy profile shows high consistency with Bonsai’s DRUID method (Pearson’s correlation coefficients: 0.983, 0.975, 0.966, and 0.943 for D_0_, D_1_, D_2_, and D_3_, respectively) and aligns with Bonsai’s likelihood method for D_0_ and D_1_ (Pearson’s correlations: 0.981 and 0.898, respectively, Supplementary Fig. 5). Notably, SNPBag avoids complex phasing or IBD calculations, reducing runtime to <0.02 seconds for a single pair of individuals, highlighting its high efficiency for large-scale applications.

## Discussion

To our knowledge, SNPBag is the first foundation model pre-trained on 1 million human whole-genome datasets. By learning from approximately 6 million genome-wide SNPs per individual, the model captures the distributions of these SNPs, their interdependencies, and the underlying haplotype structures. This endows it with superior capabilities across tasks of diverse natures: genotype imputation, haplotype phasing, population classification, and distant relatedness inference. Notably, all these capabilities were achieved using modest computational resources—8 NVIDIA A800 GPUs for whole genome pre-training and a single NVIDIA A800 or RTX3090 GPU for inference. This highlights the efficiency of the model framework and enhances its accessibility to a broader community of genomic researchers and developers. SNPBag represents a pioneering step toward a unified foundation model framework for global human genome repositories, establishing a viable new paradigm for digesting the rapidly expanding volume of genomic data under a universal AI framework. Future improvements will likely come from incorporating additional types of genetic variation, covering a greater number of loci with lower allele frequencies and novel mutations, and extending functionality to support comprehensive analyses such as sequencing data analysis, genomic annotation and admixture mapping, etc.

SNPBag operates through a fundamentally distinct mechanism compared with traditional approaches. Pre-training the base model on global genomic data enables it to learn inter-site dependencies, haplotype architectures, and even cryptic demographic structures inherent in the training data. In essence, base model pre-training involves learning a condensed, reformulated representation of the human genomic diversity— capturing both its breadth and fine-grained details—analogous to how large language models (LLMs) internalize patterns from vast text corpora. Fine-tuning for specific tasks primarily involves extracting pre-existing knowledge or patterns through representational realignment and refinement. The model inference reduces to layer-wise propagation, guided by learned weights, generating outputs based on the stored genomic knowledge.

This work carries several important implications. First, in principle, the base model can leverage datasets of virtually unbounded scale across global biobanks. Scaling laws suggest that performance will continue to improve as data scale increases, indicating that foundation-model-based inference may ultimately surpass all traditional methods when trained on sufficiently large empirical datasets, paralleling the trajectory of LLMs. Second, inference with foundation models—centered on retrieving pre-learned knowledge and probability distributions and operating with linear computational complexity—is inherently more efficient than traditional statistical methods, which typically require intensive *ab initio* computations and typically exhibit non-linear complexity. This distinction accounts for the substantial differences in inference latency between SNPBag and traditional tools for genotype imputation and phasing. Third, whereas traditional methods require multi-individual-level data, such as reference panels or population cohorts, SNPBag can perform inference on a single-individual without external data, greatly improving efficiency and flexibility.

The exponential growth of personal genomic data presents both opportunities and significant challenges for modern human genomic analysis. Privacy concerns, driven by re-identification risks and stringent regulations, restrict data sharing and hinder global collaborations in large-scale studies such as GWAS, where traditional imputation and phasing rely on extensive shared reference panels. The SNPBag framework offers a potential solution: confidential training can occur within each biobank, with only model weights shared or online inference services provided for third-party applications. A “Global genome Model” could be envisioned through iterative or distilled training across different national bio-banks, ultimately enabling genomic inference service based on the most comprehensive and deep understanding of global human diversity. At the same time, SNPBag demonstrates that the compact genome embeddings (0.75MB with further compression potential) facilitate data handling, support multitask inference, and allow simpler and more secure encryption than raw genomic data.

Ultimately, the goal of large-scale genome sequencing is to predict and interpret heritable phenotypes from genomic data. GWAS studies have identified large number of loci associated with complex traits, yet each explains only a small fraction of phenotypic variance (*21, 47*). Multivariate methods such as polygenic scores (PGS), polygenic risk scores (PRS), and GTCA models aggregates SNP effects linearly, but they fail to account for linkage disequilibrium (LD) complexity and show limited predictive power due to missing heritability(*48, 49*). Neural network-based approaches have attempted to capture the non-linear interactions, but at limited variant sites (*50*). By contrast, SNPBag encodes whole-genome SNPs, LD and haplotype structures within a machine learning architecture driven by attention mechanism—one that excels at learning long-range dependencies and high-order interactions across large number of tokens. Thus, SNPBag provides an ideal framework for robust phenotype prediction by efficiently modeling high-order epistatic interactions across the genome.

## Methods

### Base model pre-training

The SNPBag base model was developed using a Bidirectional Encoder Representations from Transformers (BERT) architecture, following standard BERT pre-training protocols. Genotype sequences, coded as 0, 1, or 2 (representing the number of derived alleles), were randomly masked at 85–99% of positions (replaced with a mask token, 3) and paired with corresponding SNP identifiers to form input sequences of 81,920 tokens. Each token was projected into a 512-dimensional space via linear layers to meet the encoder block’s input requirements. The encoder block consisted 16 stacked layers of bidirectional transformer encoder, each incorporating 16 attention heads with sliding window attention. The sliding window size was set to 128 tokens for the whole-genome model and 256 tokens for the chr22 model. The encoder block output was decoded by a multilayer perceptron (MLP) to predict masked genotype values. The model was optimized using cross-entropy loss against the original unmasked genotypes.

Two base models with this architecture were pre-trained: a chromosome 22 (chr22) model and a whole-genome model. The chr22 model was pre-trained on 1 million genotype sequences simulated from 3,202 samples from the 1,000 Genomes Project (1kG) 30x high-coverage dataset. We grouped these 3,202 individuals into 5 super-populations as ancestors (Africans, Admixed Americans, East Asian, European, and South Asian). Within each super-population, we simulated the offspring of 8^th^ generation using RESHAPE software. This data, referred to as RESHAPE data, was also used in fine-tuning of genotype imputation and haplotype phasing. After pre-training of chr22 base model, its performance was evaluated on 216 out of 929 independent Human Genome Diversity Project (HGDP) samples. The whole-genome model was pre-trained on 1 million synthetic genomes from the HAPNEST project, incorporating 1kG and HGDP samples as ancestors to enhance genetic diversity. With approximately 836 million parameters, it was evaluated for genome embedding generation, population classification, and distant relatedness inference.

Structural variants were excluded, and only single nucleotide polymorphism sites shared between the 1kG and HGDP datasets with a minor allele frequency (MAF) >1% (based on 1kG) were selected, resulting in 6,030,514 sites for the whole-genome model. During training, SNPs on the same chromosome were organized into consecutive contigs ranging from 1,024 to 81,920 sites. For the chr22 benchmark model, a single contig of 81,920 sites was used, covering approximately 92% of chromosome 22. The first 6,000 sites and the last 967 sites of chromosome 22 were excluded due to the scarcity of SNP array probes.

### Genotype imputation

Imputation accuracy was evaluated using the Illumina Omni2.5 array as a representative. For chromosome 22, we analyzed 81,920 SNP sites, of which 16,230 (20%) were present on the Omni2.5 array, with the remaining 80% requiring imputation. Complete genotype sequences from 216 HGDP individuals (4 randomly selected individuals per 54 populations) were used for evaluation. Non-array sites were masked according to the Omni2.5 array design, and masked sequences were input to several methods: SNPBag base, SNPBag omni, and conventional imputation pipelines. The SNPBag base method applied the pre-trained chr22 model directly for inference. The SNPBag omni method involved fine-tuning the chr22 model with the RESHAPE data, on imputation tasks using masking patterns aligned with the Omni2.5 array. Conventional imputation followed pipelines benchmarked by De Marino et al. (*51*), combining phasing (SHAPEIT4, BEAGLE5.2, or EAGLE2, with or without the 1kG reference panel) and imputation (IMPUTE5, MINIMAC4, or BEAGLE5.2). Performance was stratified by MAF bins and continental population groupings. MAF bins were determined by collecting roughly equal numbers of sites in each MAF interval.

### Haplotype phasing

The chr22 base model was fine-tuned for haplotype phasing using 1 million phased genotype sequences synthesized via RESHAPE from 1kG samples (the RESHAPE data). With unphased genotypes as input and a new MLP as the decoder, the model was trained to predict allelic state switches (ancestral or derived) between adjacent heterozygous sites, optimized using cross-entropy loss against true switch sequences. Note that switch sequences and haplotype sequences can be translated to each other. Model performance was evaluated using switch error rates on 26 physically phased HGDP genomes, comparing SNPBag against conventional phasing tools (SHAPEIT4, BEAGLE5.2, or EAGLE2, with or without the 1kG reference panel).

### Genome embedding

The whole-genome base model was fine-tuned to generate genome embeddings. Each genome was segmented into 2,934 contigs of 2,048 SNPs each. Contig sequences were randomly masked at 75–85% of sites and processed by the base model’s encoder. The encoder’s output (2,048 × 512) of each contig was compressed into a 128-dimensional embedding using two sequential MLPs. The 1st MLP compresses sequence dimension from 2,048 to 16. The 2nd compresses model dimension from 512 to 8. This was flattened to obtain a contig embedding of 128 dimensions (16 × 8). A third MLP predicted the 2,048 SNP genotypes from the contig embedding, optimized via cross-entropy loss until achieving 95% genotype recovery accuracy. At last, contig embeddings were concatenated to form a genome embedding (2,934 × 128) for each individual.

### Population classification

A lightweight neural network was trained for population classification using genome embeddings from 3,202 1kG individuals and their population labels. The network comprised three MLPs: two sequentially reduced the 2,934 × 128 genome embeddings to a 6 × 6 representation, and a third projected this onto 19 population categories defined by Gaspar and Breen (2019). Performance was assessed via 10 iterations of 5-fold cross-validation, randomly partitioning the 3,202 individuals into training and test sets (4:1 ratio) per iteration. F1 scores and other metrics were computed on test sets and averaged across iterations. We also did the same analyses for all 26 populations present in 1kG data.

### Relatedness inference

From 2,504 unrelated individuals of 1kG phase 3 panel, 14 of 26 populations were selected for their low levels of admixture. Each population was split into training and test sets, with each test set fixed at 20 individuals. Using ped-sim software, we simulated 40 pedigree trees for training and 4 for testing per population. Each tree, spanning 11 generations with 40 individuals, included 20 1kG genomes and 20 synthetic genomes, capturing pairwise relatedness from degree 1 to 19 and unrelated pairs (Supplementary Fig. 6). This yielded 12,320 synthetic genomes, 759,327 training pairs, and 73,345 test pairs. Genome embeddings were generated using SNPBag, and pairs of embeddings were combined for relatedness classification. A lightweight neural network with MLPs for dimensionality reduction and classification was trained for this task.

## Acknowledgement

The research was supported by the Project of Sanya Yazhou Bay Science and Technology City, Grant No: SKJC-2024-02-002. It was also supported by High-performance Computing Platform of YZBSTCACC (YaZhou Bay Science and Technology City Advanced Computing Center). We thank Prof. Haipeng Li and Prof. Shuhua Xu for constructive discusssion.

## Declaration of generative AI and AI-assisted technologies in the writing process

During the preparation of this work, the authors used GPT-5 in order to facilitate the process of proofreading the contents of our draft. After using this tool/service, the authors reviewed and edited the content as needed and take full responsibility for the content of the publication (*52*).

## References

1. J. Jumper et al., Highly accurate protein structure prediction with AlphaFold. Nature 596, 583–589 (2021).

2. F. Yang et al., scBERT as a large-scale pretrained deep language model for cell type annotation of single-cell RNA-seq data. Nat Mach Intell 4, 852–866 (2022).

3. C. V. Theodoris et al., Transfer learning enables predictions in network biology. Nature 618, 616–624 (2023).

4. H. Cui et al., scGPT: toward building a foundation model for single-cell multi-omics using generative AI. Nat Methods 21, 1470–1480 (2024).

5. H. Dalla-Torre et al., Nucleotide Transformer: building and evaluating robust foundation models for human genomics. Nat Methods, doi: 10.1038/s41592-024-02523-z (2024).

6. E. Nguyen et al., Sequence modeling and design from molecular to genome scale with Evo. Science 386, eado9336 (2024).

7. The International HapMap Consortium, The International HapMap Project. Nature 426, 789–796 (2003).

8. International Human Genome Sequencing Consortium et al., Initial sequencing and analysis of the human genome. Nature 409, 860–921 (2001).

9. O. G. Bahcall, UK Biobank — a new era in genomic medicine. Nat Rev Genet 19, 737–737 (2018).

10. The All of Us Research Program Genomics Investigators et al., Genomic data in the All of Us Research Program. Nature 627, 340–346 (2024).

11. Z. Chen et al., China Kadoorie Biobank of 0.5 million people: survey methods, baseline characteristics and long-term follow-up. International Journal of Epidemiology 40, 1652–1666 (2011).

12. T. Tanaka et al., Integrating biomedical and clinical data with BioBank Japan. Nat Cardiovasc Res 1, 597–598 (2022).

13. R. Chadwick, The Icelandic database---do modern times need modern sagas? BMJ 319, 441–444 (1999).

14. L. Leitsalu et al., Cohort Profile: Estonian Biobank of the Estonian Genome Center, University of Tartu. Int. J. Epidemiol. 44, 1137–1147 (2015).

15. M. I. Kurki et al., FinnGen provides genetic insights from a well-phenotyped isolated population. Nature 613, 508–518 (2023).

16. S. Scholtens et al., Cohort Profile: LifeLines, a three-generation cohort study and biobank. Int J Epidemiol 44, 1172–1180 (2015).

17. W. Liu et al., The 1% gift to humanity: The Human Genome Project II. Cell Res 34, 747–750 (2024).

18. OpenAI et al., GPT-4 Technical Report. arXiv arXiv:2303.08774 [Preprint] (2024). 10.48550/arXiv.2303.08774.

19. D. Guo et al., DeepSeek-R1 incentivizes reasoning in LLMs through reinforcement learning. Nature 645, 633–638 (2025).

20. The 1000 Genomes Project Consortium et al., A global reference for human genetic variation. Nature 526, 68–74 (2015).

21. E. Sollis et al., The NHGRI-EBI GWAS Catalog: knowledgebase and deposition resource. Nucleic Acids Research 51, D977–D985 (2023).

22. C. Fuchsberger et al., minimac2: faster genotype imputation. Bioinformatics 31, 782–784 (2015).

23. P.-R. Loh et al., Reference-based phasing using the Haplotype Reference Consortium panel. Nat Genet 48, 1443–1448 (2016).

24. S. Rubinacci et al., Genotype imputation using the Positional Burrows Wheeler Transform. PLOS Genetics 16, e1009049 (2020).

25. B. L. Browning et al., Fast two-stage phasing of large-scale sequence data. Am J Hum Genet 108, 1880–1890 (2021).

26. R. J. Hofmeister et al., Accurate rare variant phasing of whole-genome and whole-exome sequencing data in the UK Biobank. Nat Genet 55, 1243–1249 (2023).

27. S. Purcell et al., PLINK: a tool set for whole-genome association and population-based linkage analyses. Am J Hum Genet 81, 559–575 (2007).

28. A. Diaz-Papkovich et al., UMAP reveals cryptic population structure and phenotype heterogeneity in large genomic cohorts. PLOS Genetics 15, e1008432 (2019).

29. H. A. Gaspar, G. Breen, Probabilistic ancestry maps: a method to assess and visualize population substructures in genetics. BMC Bioinformatics 20, 116 (2019).

30. C. D. Huff et al., Maximum-likelihood estimation of recent shared ancestry (ERSA). Genome Res. 21, 768–774 (2011).

31. M. D. Ramstetter et al., Benchmarking Relatedness Inference Methods with Genome-Wide Data from Thousands of Relatives. Genetics 207, 75–82 (2017).

32. E. M. Jewett et al., Bonsai: An efficient method for inferring large human pedigrees from genotype data. The American Journal of Human Genetics 108, 2052–2070 (2021).

33. J. Chen, X. Shi, Sparse Convolutional Denoising Autoencoders for Genotype Imputation. Genes (Basel) 10, 652 (2019).

34. K. Kojima et al., A genotype imputation method for de-identified haplotype reference information by using recurrent neural network. PLOS Computational Biology 16, e1008207 (2020).

35. T. Naito et al., A deep learning method for HLA imputation and trans-ethnic MHC fine-mapping of type 1 diabetes. Nat Commun 12, 1639 (2021).

36. R. Dias et al., Rapid, Reference-Free human genotype imputation with denoising autoencoders. Elife 11, e75600 (2022).

37. M. Song et al., An autoencoder-based deep learning method for genotype imputation. Front. Artif. Intell. 5 (2022).

38. M. E. Mowlaei et al., Split-Transformer Impute (STI): Genotype Imputation Using a Transformer-Based Model. bioRxiv [Preprint] (2023). 10.1101/2023.03.05.531190.

39. X. Huang et al., Harnessing deep learning for population genetic inference. Nat Rev Genet 25, 61–78 (2024).

40. T. Cavinato et al., A resampling-based approach to share reference panels. Nat Comput Sci 4, 360–366 (2024).

41. S. Wharrie et al., HAPNEST: efficient, large-scale generation and evaluation of synthetic datasets for genotypes and phenotypes. Bioinformatics 39, btad535 (2023).

42. O. Delaneau et al., Accurate, scalable and integrative haplotype estimation. Nat Commun 10, 5436 (2019).

43. M. W. Snyder et al., Haplotype-resolved genome sequencing: experimental methods and applications. Nat Rev Genet 16, 344–358 (2015).

44. S. E. Aalbers et al., Analyzing population structure for forensic STR markers in next generation sequencing data. Forensic Science International: Genetics 49, 102364 (2020).

45. M. C. Lancaster et al., Detection of distant relatedness in biobanks to identify undiagnosed cases of Mendelian disease as applied to Long QT syndrome. Nat Commun 15, 7507 (2024).

46. A. Gusev et al., Whole population, genome-wide mapping of hidden relatedness. Genome Res 19, 318–326 (2009).

47. R. J. F. Loos, 15 years of genome-wide association studies and no signs of slowing down. Nat Commun 11, 5900 (2020).

48. A. Abdellaoui et al., 15 years of GWAS discovery: Realizing the promise. The American Journal of Human Genetics 110, 179–194 (2023).

49. X. Yang et al., Polygenic scores in cancer. Nat Rev Cancer 23, 619–630 (2023).

50. A. I. Sigurdsson et al., Deep integrative models for large-scale human genomics. Nucleic Acids Res 51, e67 (2023).

51. A. D. Marino et al., A comparative analysis of current phasing and imputation software. PLOS ONE 17, e0260177 (2022).

52. Y. Bai et al., How our authors are using AI tools in manuscript writing. Patterns 5, 101075 (2024).

