## Supplementary information for "Towards a universal foundation model for biobank-scale human genome variation"

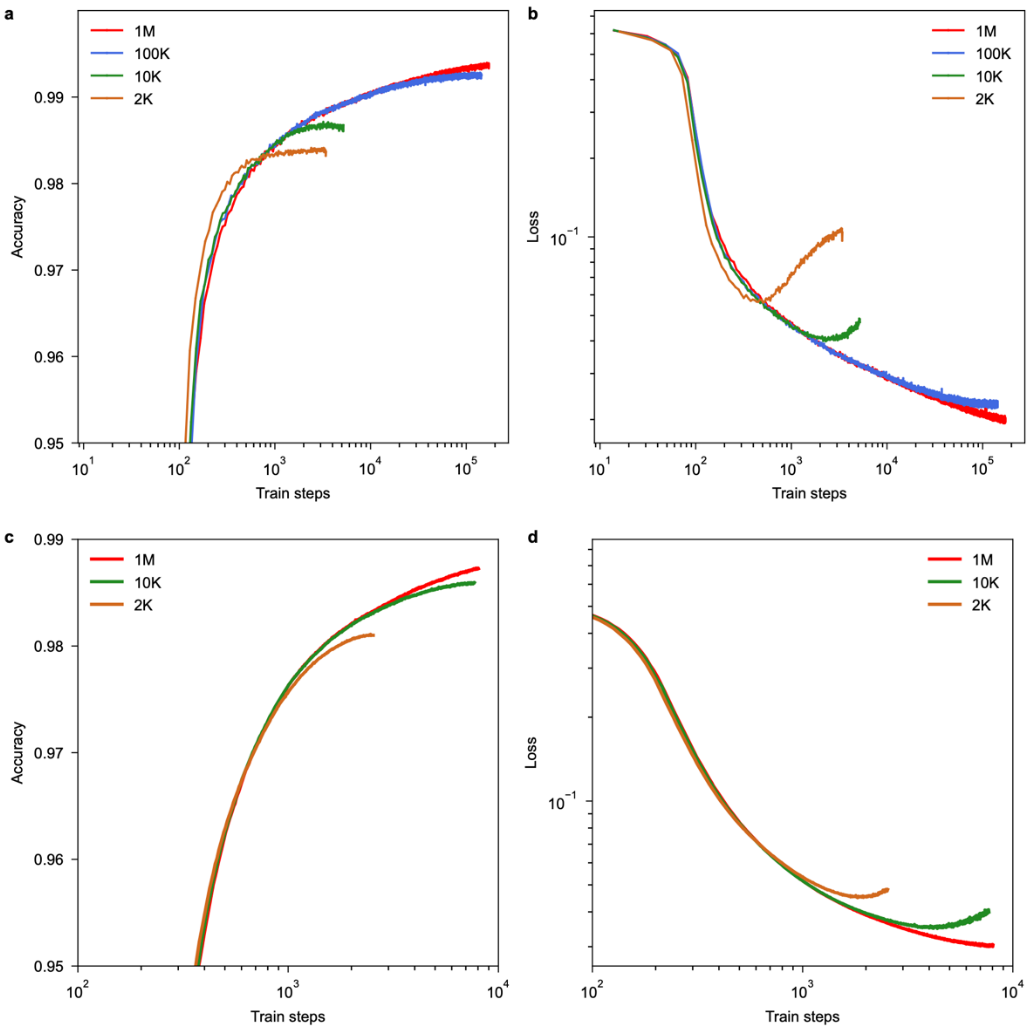


**Supplementary Fig. 1 | Scaling tests for fine-tuning of genotype imputation (a,b) and haplotype phasing (c,d). a,b,** validation accuracy curves (**a**) and loss curves (**b**) for SNPBag during fine-tuning of genotype imputation, across different training sample sizes: 1 million (red), 100K (blue), 10K (green) and 2K (brown). **c,d,** validation accuracy curves (**c**) and loss curves (**d**) for SNPBag during fine-tuning of haplotype phasing, across different training sample sizes: 1 million (red), 10K (green) and 2K (brown).

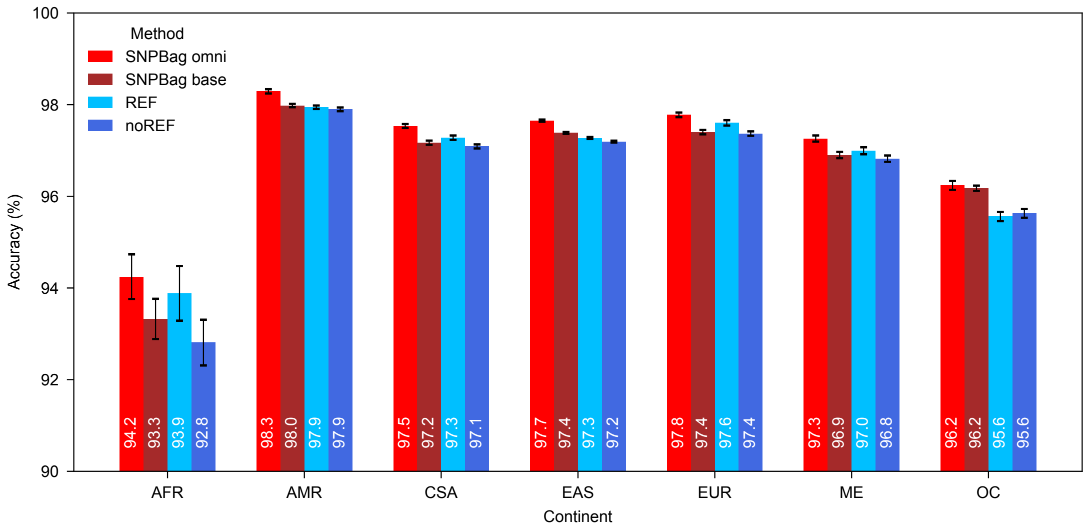


**Supplementary Fig. 2 | Imputation accuracy of different methods across continental groups.** Conventional methods with and without reference panel were averaged as REF and noREF. Continent groups were Africans (AFR), Americans (AMR), Central and South Asians (CSA), East Asians (EAS), Europeans (EUR), Middle East populations (ME) and Oceanians (OC). Error bars represent the standard errors of mean estimations.

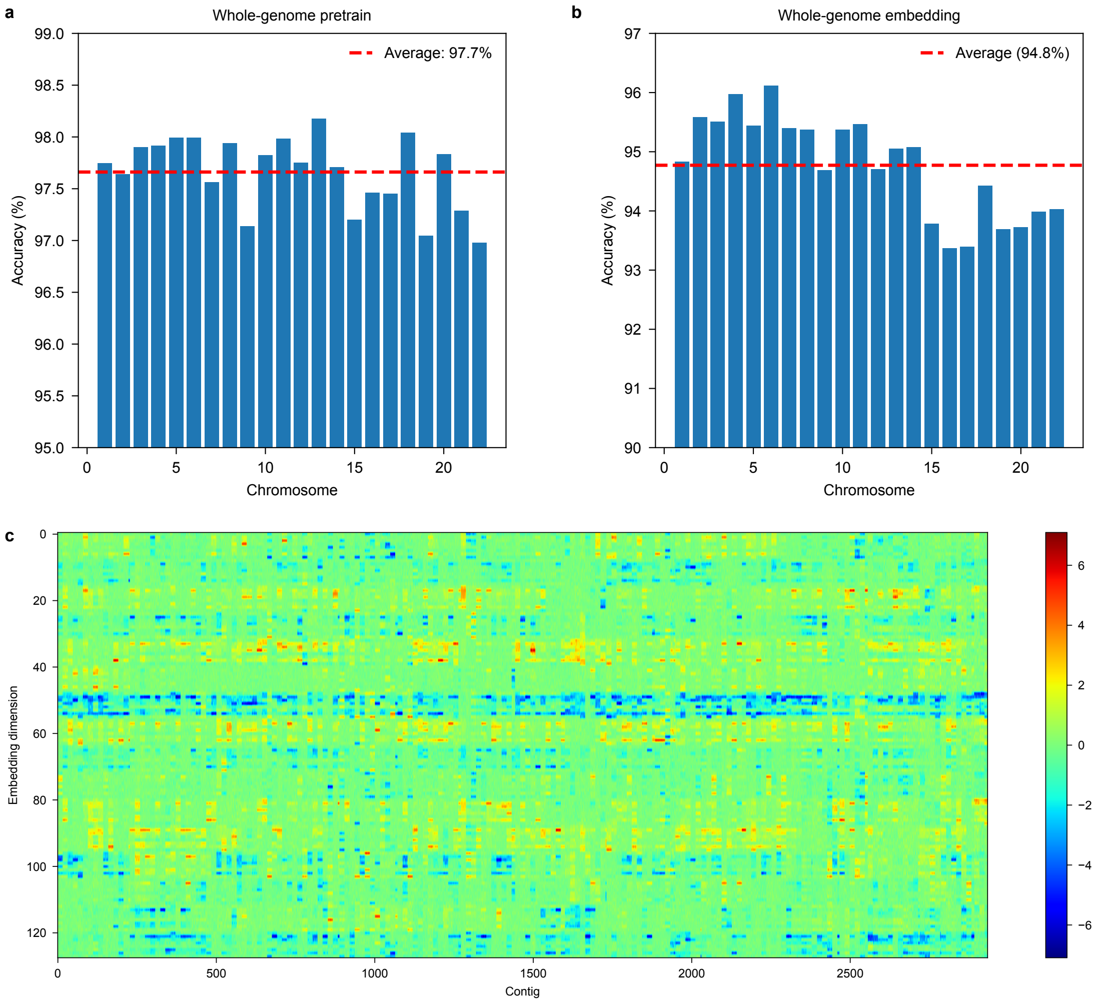


**Supplementary Fig. 3 | Embeddings of whole-genome SNPs**. **a,** whole-genome pre-training quality across 22 chromosomes, evaluated via accuracy on OMNI2.5 array genotype imputation in 216 HGDP samples. Red horizontal line represents the average across chromosomes. **b,** whole-genome embedding quality, assessed by genotype recovery accuracy for 75–85% masked genotypes. **c,** graphical representation of SNPBag's whole-genome embedding for 1 individual.

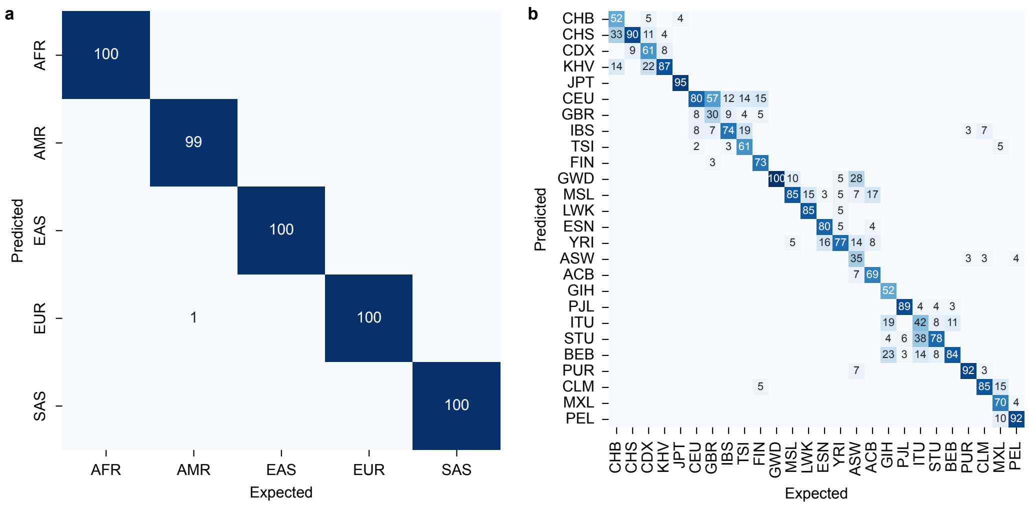


**Supplementary Fig. 4 | Performance of population classification. a,** confusion matrix showing the classification results for the 5 continental populations, with values shown as percentages; **b,** confusion matrix showing the classification results for the 26 populations, with values shown as percentages of individuals with expected classes.

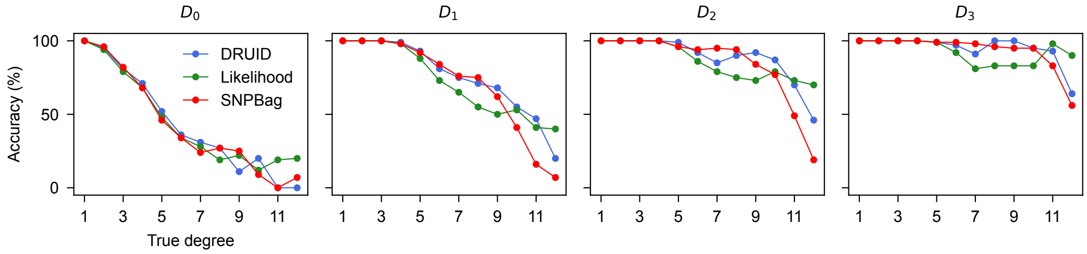


**Supplementary Fig. 5 | Performance comparison of the relatedness estimation.** Results of DRUID (blue) and Likelihood (green) are from 2 algorithms in package Bonsai. Four levels of error tolerance: exact match ($D_0$), up to 1-degree error ($D_1$), up to 2-degree error ($D_2$), and up to 3-degree error ($D_3$). Pearson correlation coefficient between SNPBag and DRUID, and between SNPBag and the Likelihood algorithm, are as follows: $D_0$ (0.983 and 0.981), $D_1$ (0.975 and 0.898), $D_2$ (0.966 and 0.689), and $D_3$ (0.943 and 0.143), respectively.

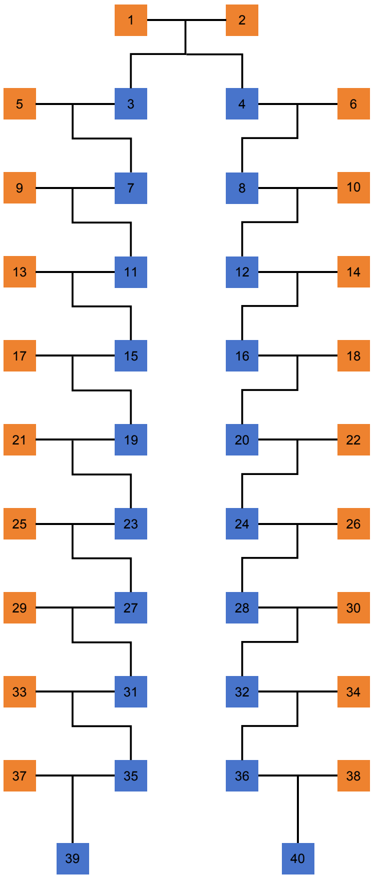


**Supplementary Fig. 6 | Simulation of pedigree.** Pairs of individuals with relatedness from 1 to 19 degrees were created by simulating pedigrees with software ped-sim. Each simulated pedigree tree, spanning 11 generations with 40 individuals, included 20 1kG genomes (orange) and 20 synthetic genomes (blue). Numbers represent sample ID in each tree. Sex chromosomes were excluded. Genders were ignored.
